# Ciliary Hedgehog signaling regulates cell survival to build the facial midline

**DOI:** 10.1101/2021.03.18.436057

**Authors:** Shaun Abrams, Jeremy F. Reiter

## Abstract

Craniofacial defects are among the most common phenotypes caused by ciliopathies, yet the developmental and molecular etiology of these defects is poorly understood. We investigated multiple mouse models of human ciliopathies (including *Tctn2, Cc2d2a* and *Tmem231* mutants) and discovered that each displays hypotelorism, a narrowing of the midface. As early in development as the end of gastrulation, *Tctn2* mutants displayed reduced activation of the Hedgehog (HH) pathway in the prechordal plate, the head organizer. This prechordal plate defect preceded a reduction of HH pathway activation and *Shh* expression in the adjacent neurectoderm. Concomitant with the reduction of HH pathway activity, *Tctn2* mutants exhibited increased cell death in the neurectoderm and facial ectoderm, culminating in a collapse of the facial midline. Enhancing HH signaling by decreasing the gene dosage of a negative regulator of the pathway, *Ptch1,* decreased cell death and rescued the midface defect in both *Tctn2* and *Cc2d2a* mutants. These results reveal that ciliary HH signaling mediates communication between the prechordal plate and the neurectoderm to provide cellular survival cues essential for development of the facial midline.

## Introduction

Primary cilia are microtubule-based organelles present on diverse vertebrate cell types and critical for development. Primary cilia function as specialized cellular signaling organelles that coordinate multiple signaling pathways, including the Hedgehog pathway (Zaghloul & Brugmann, 2011). Defects in the structure or signaling functions of cilia cause a group of human syndromes, collectively referred to as ciliopathies, which can manifest in diverse phenotypes including cystic kidneys, retinal degeneration, cognitive impairment, respiratory defects, left-right patterning defects, polydactyly, and skeletal defects (Baker & Beales, 2009; Schwartz, Hildebrandt, Benzing, & Katsanis, 2011; Tobin & Beales, 2009). In addition to these phenotypes, craniofacial defects including cleft lip/palate, high-arched palate, jaw disorders, midface dysplasia, craniosynostosis, tongue abnormalities, abnormal dentition and tooth number and exencephaly, are observed in approximately one-third of individuals with ciliopathies (Brugmann, Cordero, & Helms, 2010b; Zaghloul & Brugmann, 2011). The molecular and developmental etiology of these craniofacial abnormalities remains poorly understood.

HH signaling is intimately involved in forebrain and midface development (D. H. A. J. A. Helms, 1999; Rubenstein & Beachy, 1998). In humans, inherited mutations that compromise pathway activity impair forebrain development and cause hypotelorism (Fuccillo, Joyner, & Fishell, 2006; D. H. A. J. A. Helms, 1999; Hu & Marcucio, 2008; Marcucio, Cordero, Hu, & Helms, 2005; Muenke & Beachy, 2000; Young, Chong, Hu, Hallgrimsson, & Marcucio, 2010). For example, mutations in *SHH* lead to holoprosencephaly (Chiang et al., 1996; Cohen & Shiota, 2002). Meckel syndrome (MKS), a severe ciliopathy, is also characterized by holoprosencephaly and hypotelorism (Ben Chih et al., 2011; Dowdle et al., 2011; Garcia-Gonzalo et al., 2011). MKS-associated genes encode proteins that form a complex that comprises part of the transition zone, a region of the ciliary base that regulates ciliogenesis and ciliary membrane protein composition in a tissue-specific manner (Ben Chih et al., 2011; Dowdle et al., 2011; Garcia-Gonzalo et al., 2011; Roberson et al., 2015). Disruption of this transition zone complex results in impaired the ciliary localization of several membrane-associated signaling proteins including Smoothened (SMO), Adenylyl cyclase 3 (ADCY3), Polycystin 2 (PKD2) and ARL13B (Ben Chih et al., 2011; Garcia-Gonzalo et al., 2011; Roberson et al., 2015).

We investigated the molecular underpinnings of forebrain and midface defects in ciliopathies utilizing multiple mouse mutants affecting the transition zone. The mutants exhibited forebrain and midface defects by E9.5, which persisted throughout development. In these mutants, the prechordal plate, an organizer of anterior head development, displayed defects in HH pathway activation at E8.0. These early prechordal plate defects attenuated *Shh* expression in the adjacent ventral forebrain. Decreased HH signaling increased apoptosis in the ventral neurectoderm and facial ectoderm. Surprisingly, reducing *Ptch1* gene dosage rescued the apoptosis and its corresponding midface defect. Thus, investigating the function of the transition zone has revealed a key role of prechordal plate-activated HH signaling in forebrain and midface cell survival. Moreover, our genetic results reveal that inhibition of PTCH1 can prevent ciliopathy-associated midface defects. Based on these mouse genetic models, we propose that the etiology of hypotelorism in human ciliopathies is a failure of the prechordal plate to induce SHH expression in the overlying ventral neuroectoderm.

## Results

### The ciliary MKS transition zone complex is essential for midline facial development

Individuals affected by developmental ciliopathies, such as Meckel, Orofaciodigital and Joubert syndromes, often display craniofacial phenotypes including holoprosencephaly and hypotelorism (Baker & Beales, 2009; Dowdle et al., 2011; Garcia-Gonzalo et al., 2011). To explore the etiology of these craniofacial defects, we examined the craniofacial development in *Tctn2* mouse mutants (Garcia-Gonzalo et al., 2011; Shaheen et al., 2011). TCTN2 is a component of the MKS transition zone complex critical for ciliary localization of several ciliary membrane proteins, including SMO, a key ciliary mediator of HH signaling (Ben Chih et al., 2011; Corbit et al., 2005; Garcia-Gonzalo et al., 2011) Mutations in human *TCTN2* cause Meckel and Joubert syndromes (Huppke et al., 2015; Sang et al., 2011; Shaheen et al., 2011).

*Tctn2*^+/-^ embryos were phenotypically indistinguishable from *Tctn2^+/+^* embryos (**Supplemental Figure 1)**. In contrast, embryonic day (E) 10.5 *Tctn2*^-/-^ embryos exhibited an approximately 50% decrease in infranasal distance (the distance between the nasal pits, the nostril anlage) (**Figure 1A**). One day later in gestation (E11.5), *Tctn2*^-/-^ embryos also exhibited midfacial narrowing, including hypoplasia of the frontonasal prominence and fusion of the two maxillary prominences at the midline (**Figure 1A**). Thus, TCTN2 is essential for development of the facial midline.

**Figure 1.**
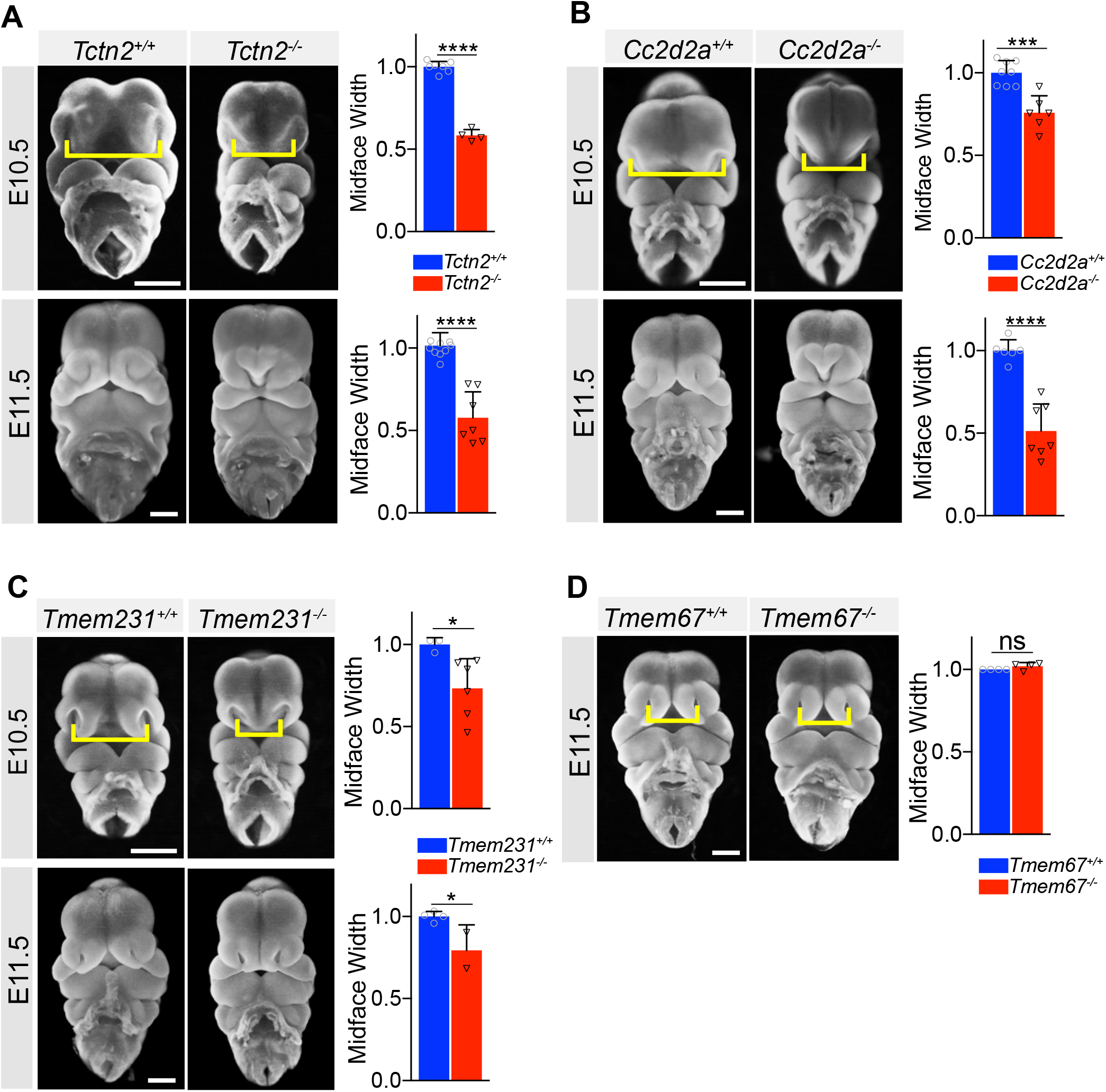
The ciliary MKS transition zone complex is essential for midline facial development. Frontal view images of *Tctn2* (A), *Cc2d2a* (B), *Tmem231* (C) wildtype and null embryos at E10.5 and E11.5. *Tmem67* null embryos (D) display normal midface width at E11.5.Quantification of midface width (denoted by yellow brackets) at respective timepoints was measured via one-way Anova followed by Tukey’s multiple comparisons test. Data are expressed as mean and error bars represent the SD with individual data points. Scale bar indicates 500μm. p<0.05 (*), p<0.0002 (***), p<0.0001 (****)

To assess whether this narrowing of the facial midline is specific to TCTN2, we examined the possible involvement of two other components of the MKS complex, TMEM231 and CC2D2A, in craniofacial development. Human *CC2D2A* mutations cause Meckel and Joubert syndromes, and *TMEM231* mutations cause Meckel, Joubert and Orofaciodigital syndromes (Noor et al., 2008; Roberson et al., 2015; Shaheen, Ansari, Mardawi, Alshammari, & Alkuraya, 2013; Srour et al., 2012; Tallila, Jakkula, Peltonen, Salonen, & Kestilä, 2008). Similar to *Tctn2* mutants, both E10.5 *Cc2d2a* and *Tmem231* mutant embryos exhibited decreased infranasal distance (**Figure 1B and Figure 1C**, respectively). The similarity of the midline hypoplasia in all three transition zone mutants suggested a common etiology.

We also examined the involvement of a fourth member of the MKS complex, TMEM67, in craniofacial development. Human mutations in *TMEM67* also cause Meckel and Joubert syndromes (Otto et al., 2009; Smith et al., 2006). Mutation of mouse *Tmem67* causes phenotypes that are less severe than *Tctn2*, *Tmem231* or *Cc2d2a* (Garcia-Gonzalo et al., 2011). The mild phenotype of *Tmem67* mutants may be attributable to its dispensability for ciliary accumulation of HH pathway activator SMO (Garcia-Gonzalo et al., 2011). Unlike *Tctn2*, *Tmem231* and *Cc2d2a* mutants, *Tmem67* mutants did not exhibit altered infranasal distance (**Figure 1D**). Thus, some, but not all, MKS components are critical for early facial midline development.

Given the central role of the neural crest in craniofacial development as the main source of craniofacial mesenchyme (Santagati & Rijli, 2003), we tested whether transition zone disruption in this tissue is the origin of the midline defect seen in *Tctn2* mutants. Interestingly, conditional deletion of *Tctn2* in the neural crest using the *Wnt1-Cre* driver did not result in hypotelorism (**Supplemental Figure 2**). This result suggests that the neural crest is not the origin of the midline phenotype and led us to investigate a role for *Tctn2* in the prechordal plate, an early organizing center critical for development of anterior head structures.

### *Tctn2* mutants exhibit defects in prechordal plate differentiation soon after gastrulation

As *Tctn2*, *Cc2d2a* and *Tmem231* mutants all displayed facial midline defects at midgestation, we hypothesized that they shared a role in a patterning event early in craniofacial development. One organizing center critical for early forebrain and craniofacial development is the prechordal plate (Camus et al., 2000; Kiecker & Niehrs, 2001; Muenke & Beachy, 2000; Rubenstein & Beachy, 1998; Som, Streit, & Naidich, 2014). The prechordal plate is the anterior-most midline mesendoderm, immediately anterior to the notochord and in contact with the overlying ectoderm. Previous work demonstrated that surgical removal of the rat prechordal plate results in midface defects (Aoto et al., 2009) that seemed similar to those of the mouse *Tctn2*, *Cc2d2a* and *Tmem231* mutants.

Therefore, we analyzed the prechordal plate of *Tctn2* mutants by examining the expression of axial mesendodermal markers *T, Gsc* and *Shh*. *Gsc* is expressed specifically in the prechordal mesoderm, while *Shh* and *T* are expressed in both the prechordal mesoderm and notochord (Dale et al., 1997; Herrmann, 1991; Schulte-Merker et al., 1994). *In situ* hybridization analysis revealed that in *Tctn2* mutants, *Shh* and *T* expression in the prechordal plate and notochord were unaffected (**Figure 2A**). Therefore, TCTN2 is not essential for prechordal plate specification. In contrast, *Tctn2* mutants exhibited abrogated *Gsc* expression in the prechordal plate (**Figure 2B**), indicating that TCTN2 is critical for prechordal plate differentiation.

**Figure 2:**
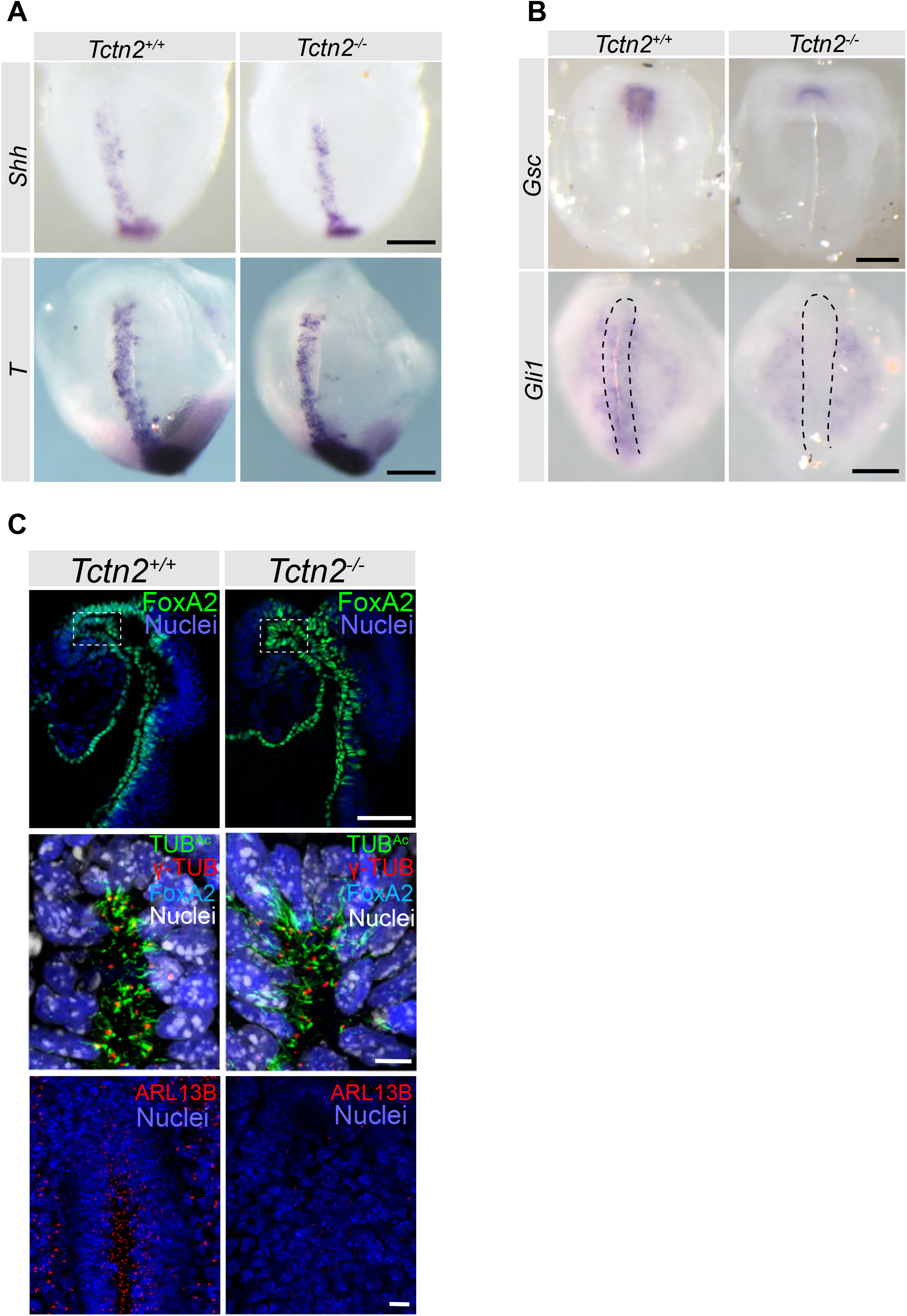
*Tctn2* mutants exhibit defects in prechordal plate differentiation soon after gastrulation. (A): WM-ISH of E8.0 embryos for axial mesendoderm markers *Shh* and *T.* (B) WM-ISH of E8.0 embryos for prechordal plate marker *Gsc* and HH pathway target *Gli*1. (C) Whole mount immunofluorecence staining for the cilia marker acetylated tubulin (TUB^Ac^), basal body marker gamma tubulin (γ-TUB) and ciliary membrane protein ARL13B in E8.0 embryos. Middle panel in C is magnified region in top panel highlighted by dotted rectangle and rotated 90 degrees. Scale bar in A-B indicates 0.2mm, C (top panel) is 100μm, C (middle and bottom panels) is 10μm

The transition zone is critical for HH signaling, and one HH protein, SHH, is essential for *Gsc* expression in the prechordal plate (Aoto et al., 2009; Ben Chih et al., 2011; Garcia-Gonzalo et al., 2011). Therefore, we investigated whether TCTN2 is required for HH signaling in the prechordal plate by examining the expression of the transcriptional target *Gli1. Tctn2* mutants exhibited reduced *Gli1* expression throughout the axial mesendoderm, including the prechordal plate (**Figure 2B**). These results indicate that TCTN2 is dispensable for the formation of the prechordal plate, but is required for midline signaling by SHH to induce *Gsc* expression in this organizing center.

TCTN2 and other members of the MKS complex are required for proper cilia formation in some tissues but not in others (Garcia-Gonzalo et al., 2011). Therefore, we examined whether TCTN2 is required for ciliogenesis in the prechordal plate. The prechordal plate expresses FOXA2 (Jin, Harpal, Ang, & Rossant, 2001), (**Figure 2C**). Co-immunostaining embryos for FOXA2 and acetylated tubulin (TUB^Ac^), a marker of cilia, revealed that *Tctn2* mutants did not display decreased ciliogenesis in the E8.5 prechordal plate (**Figure 2C**, middle panel).

In cell types in which the MKS complex is dispensable for ciliogenesis, like neural progenitors, it is required for localization of ARL13B to cilia (Garcia-Gonzalo et al., 2011). Therefore, we examined ARL13B localization in E8.0 control embryos and *Tctn2* mutants and discovered that ARL13B localization to prechordal plate cilia was attenuated without TCTN2 (**Figure 2C**, bottom panel). Thus, TCTN2 is not required for ciliogenesis in the prechordal plate, but does control ciliary composition.

### *Tctn2* mutants display decreased HH signaling in the ventral telencephalon

The axial mesendoderm helps pattern the overlying neurectoderm (Anderson & Stern, 2016; Rubenstein & Beachy, 1998). In the rostral embryo, the prechordal plate patterns the overlying ventral telencephalon via SHH (Chiang et al., 1996; Xavier et al., 2016). As extirpation of the prechordal plate results in decreased SHH activity in the basal telencephalon (Aoto et al., 2009; Aoto & Trainor, 2015), we investigated whether the prechordal plate defects observed in *Tctn2* mutants results in mispatterning of the ventral telencephalon. Although *Shh* expression in the notochord was unaffected in *Tctn2* mutants at E8.75, it was severely reduced in the ventral telencephalon (**Figure 3A**). This reduced expression of *Shh* in the ventral telencephalon persisted at E9.5 (**Figure 3A**).

**Figure 3.**
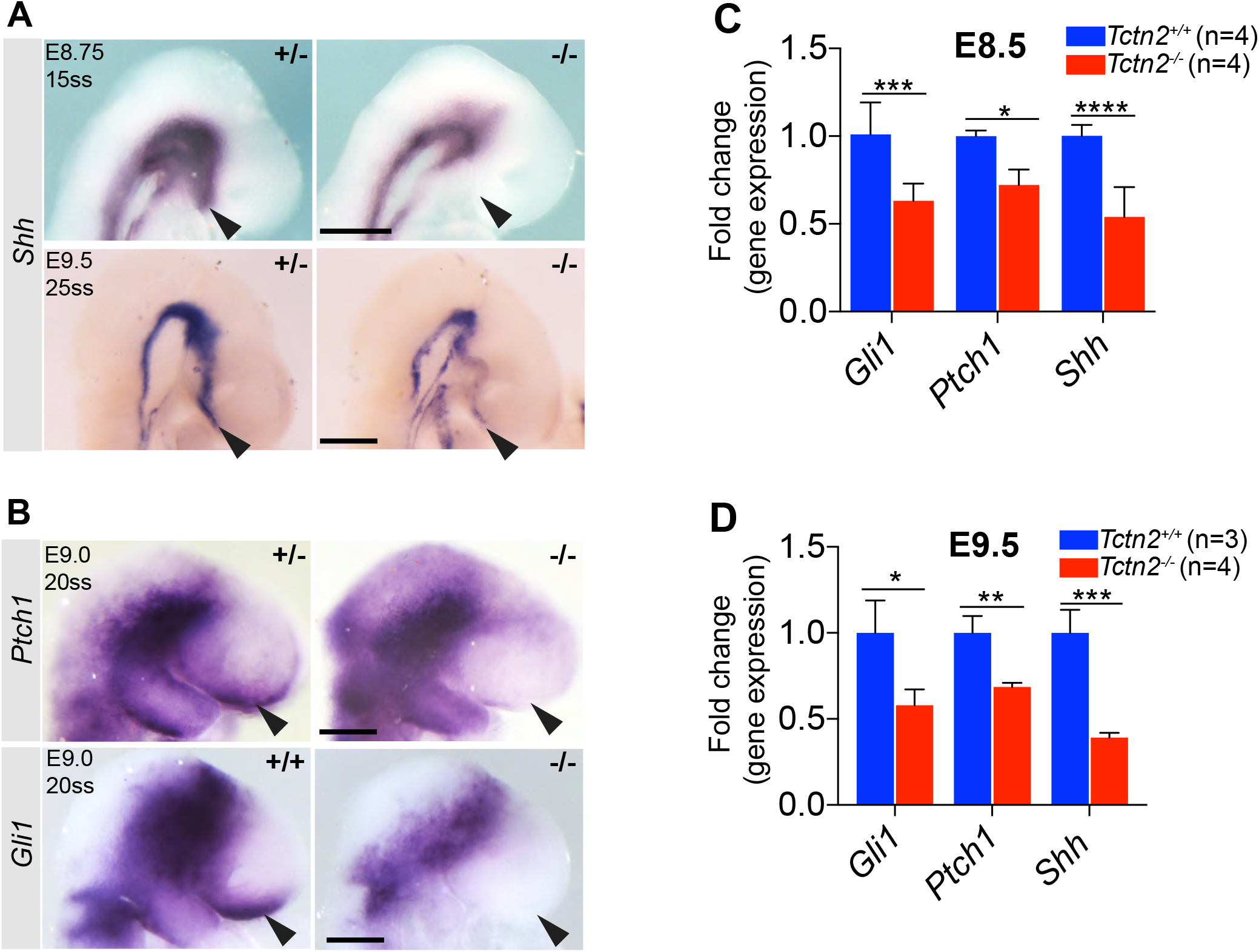
*Tctn2* mutants display decreased HH signaling in the ventral telencephalon. WM-ISH for *Shh* in *Tctn2* control and mutant embryos at E8.75 and E9.5 (A) show reduced expression in the ventral telencephalon of mutant embryos (arrowheads). WM-ISH for HH pathway targets *Ptch1* and *Gli1* also show reduced expression in the ventral telencephalon in *Tctn2* mutants (B, arrowheads). RT-qPCR analysis of RNA transcripts isolated from E8.5 and E9.5 *Tctn2* control and mutant heads (C and D respectively) show reduced levels of *Gli*, *Ptch1,* and *Shh* transcripts, consistent with WM-ISH results. Data in C, D represent mean and error bars represent the SD. Scale bar indicates 0.5mm. p<0.05 (*), p<0.001 (**) p<0.0002 (***), p<0.0001 (****)

To assess whether HH pathway activity was compromised by the absence of TCTN2, we assessed the expression of the HH transcriptional targets *Gli1* and *Ptch1*. WM-ISH of *Tctn2* mutants revealed dramatically reduced or absent expression of both *Gli1* and *Ptch1*, especially in the basal forebrain (**Figure 3B**). Consistent with the WM-ISH data, qRT-PCR analysis of E8.5 (**Figure 3C**) and E9.5 (**Figure 3D**) *Tctn2* mutant heads also revealed decreased expression of *Shh, Ptch1* and *Gli1,* revealing that *Tctn2* mutants exhibit both an early defect in prechordal plate differentiation and a defect in HH signaling in the adjacent neurectoderm.

### TCTN2 protects neurectoderm and facial ectoderm from apoptosis

In the developing craniofacial complex, SHH induces cell proliferation (D. H. A. J. A. Helms, 1999; Hu et al., 2015). Therefore, we assessed if the reduction in facial midline width in *Tctn2* mutants was due to decreased cell proliferation. More specifically, we measured cell proliferation by quantitating phospho-histone H3 in the components of the craniofacial complex – the forebrain, hindbrain, facial ectoderm and mesenchyme (**Figure 4A,B**). *Tctn2* mutants showed no differences in amount or spatial distribution of cell proliferation.

**Figure 4.**
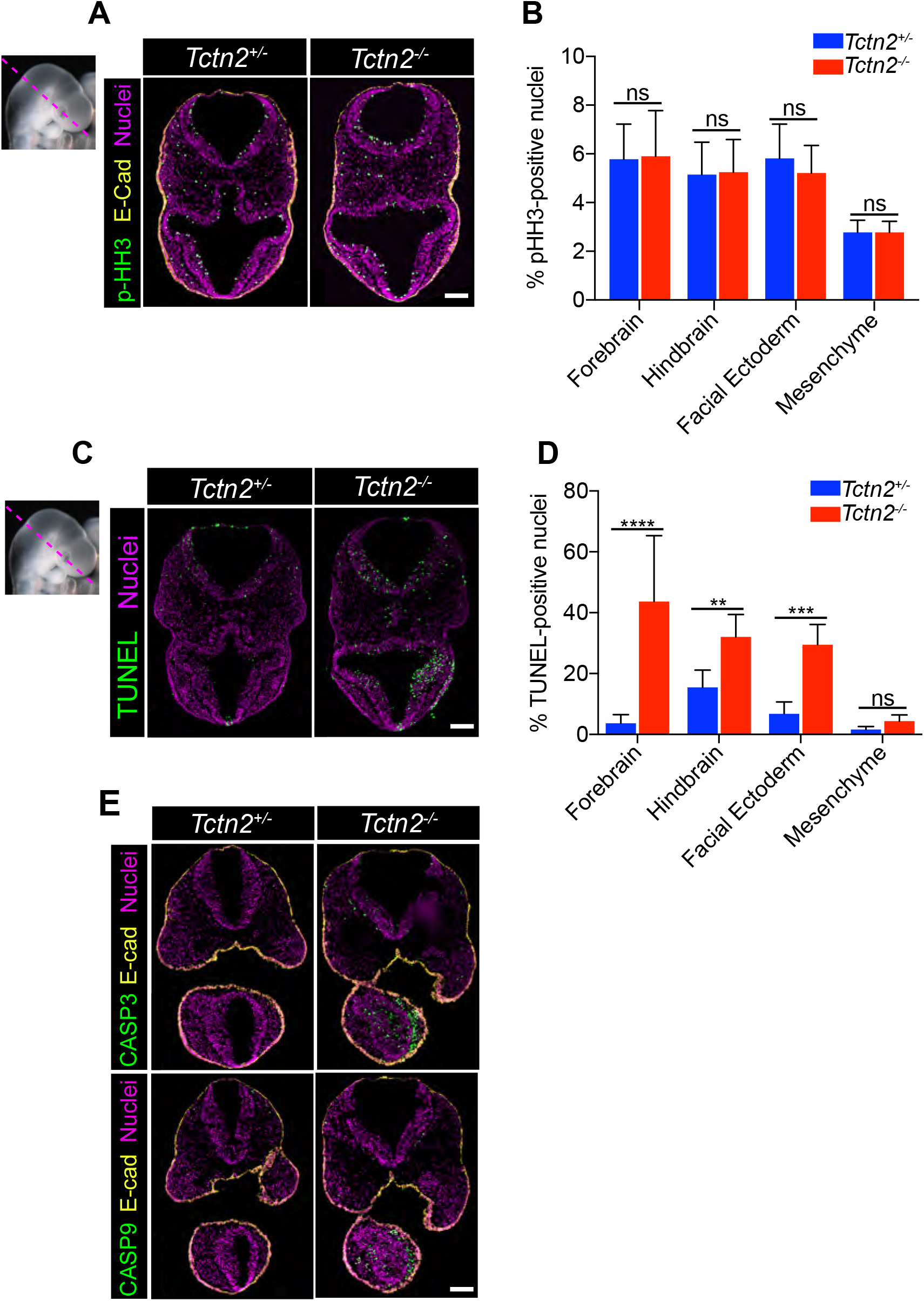
TCTN2 protects the neurectoderm and facial ectoderm from apoptosis. Immunostaining for proliferation marker phospho-histone H3 (pHH3) in transverse sections of *Tctn2* control and mutant E9.5 embryos (A) with corresponding quantification (B). (C) TUNEL staining for analysis of apoptosis in *Tctn2* E9.5 embryos with corresponding quantification (D). (E) Immunostaining of intrinsic apoptotic pathway components cleaved-caspase 3 (CASP3) and cleaved-caspase 9 (CASP9) in *Tctn2* control and mutant E9.5 embryos. Data in B, D represent the mean and errory bars represent the SD.Scale bar indicates100μm. p<0.002 (**) p<0.0002 (***), p<0.0001 (****). ns = not significant.

In other developmental contexts, HH signaling promotes cell survival (Ahlgren & Bronner-Fraser, 1999; Aoto et al., 2009; Aoto & Trainor, 2015; Litingtung & Chiang, 2000). Therefore, we assessed apoptosis in the craniofacial complex of *Tctn2* mutants. Quantification of TUNEL staining revealed that apoptosis in the mesenchyme was unaffected, but increased in the neurectoderm and facial ectoderm of *Tctn2* mutants compared to controls, and most dramatically in the ventral telencephalon (**Figure 4C,D**). To further test whether apoptosis is increased in the absence of TCTN2, we stained for activators of the intrinsic apoptotic pathway, cleaved Caspase-3 and Caspase-9 (activated CASP3 and CASP9). Both activated CASP3 and CASP9 were increased in the ventral telencephalon and facial ectoderm of *Tctn2* mutants at E9.5 (**Figure 4E**). These data indicate that TCTN2 is required to protect against cell death, but does not affect proliferation, in the neurectoderm and non-neural ectoderm. As SHH also protects neurectoderm from apoptosis (Thibert et al., 2003), we propose that TCTN2 mediates cell survival by promoting HH signaling and that the increase in cell death underlies the midfacial hypoplasia in transition zone mutants.

### Reducing *Ptch1* gene dosage rescues the facial midline defect in transition zone mutants

To assess whether decreased HH signaling is not just correlated with midfacial hypoplasia in transition zone mutants, but is causative, we investigated whether modulating the HH pathway could rescue the midface defects. We employed a strategy targeting *Ptch1*, a negative regulator of the HH pathway, by generating *Tctn2*^-/-^ *Ptch1*^-/+^ embryos and comparing them to *Tctn2*^-/-^ *Ptch1*^+/+^ embryos. Surprisingly, removing a single allele of *Ptch1* in *Tctn2* mutants restored midface width at E11.5 (**Figure 5A,B**).

**Figure 5.**
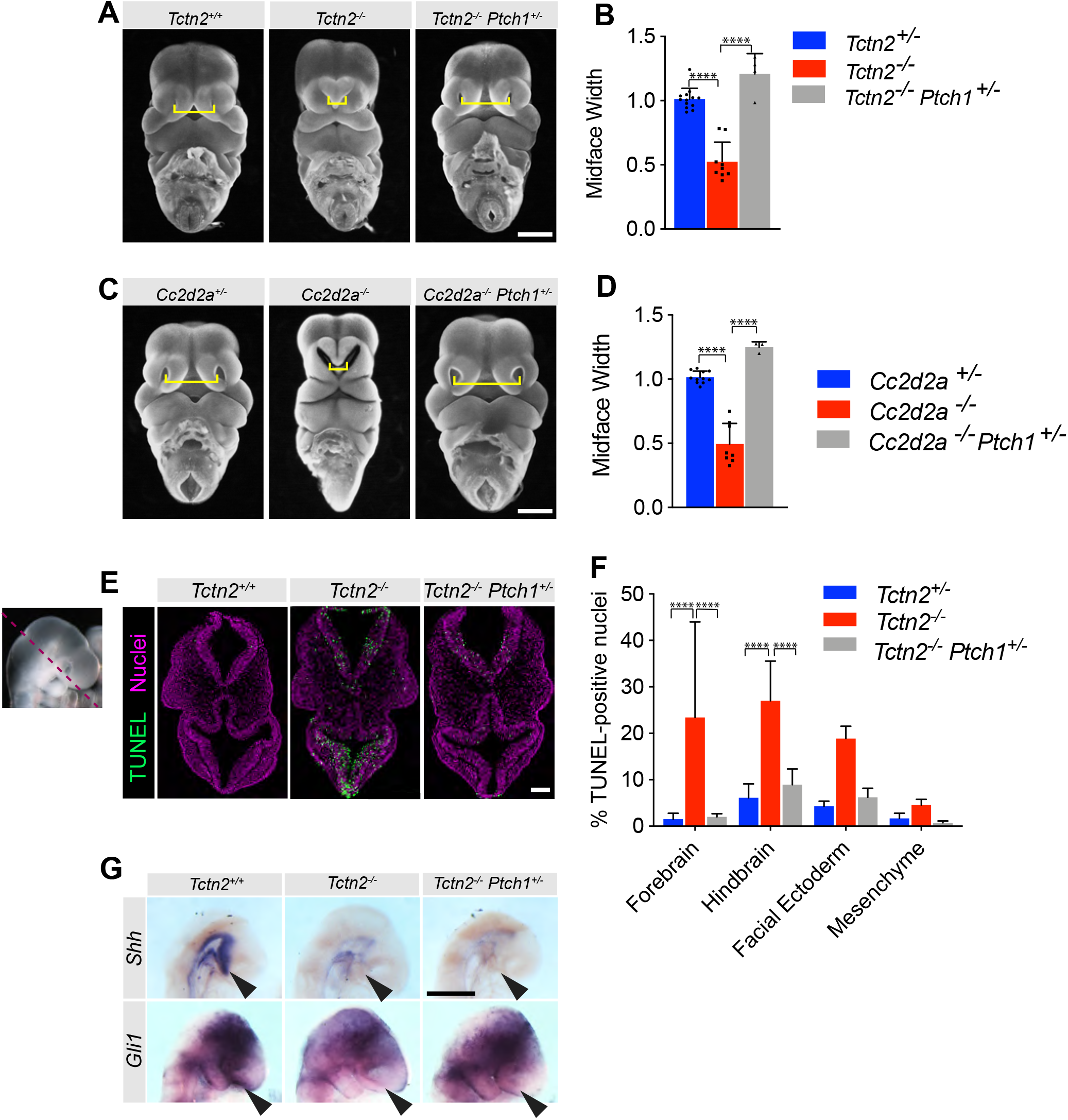
Reducing *Ptch1* gene dosage rescues the facial midline defect in transition zone mutants. (A) Frontal view images of *Tctn2* wildtype, mutant, and *Ptch1*^+/-^ rescue E11.5 embryos with corresponding midface width quantification (B). (C) Frontal view images of *Cc2d2a* wildtype, mutant, and *Ptch1*^+/-^ rescue E11.5 embryos with corresponding midface width quantification (D). (E) TUNEL assay sections of E9.5 *Tctn2* wildtype, mutant, and *Ptch1*^+/-^ rescue embryos with corresponding quantification (F). (G) WM-ISH of E9.5 *Tctn2* wildtype, mutant, and *Ptch1*^+/-^ rescue embryos for *Shh* and *Gli1*. Data in B, D, and F represent the mean and error bars represents SD. For statistical analysis one-way ANOVA was performed with Tukey’s multiple comparison test. p<0.0001 (****). Scale bars indicate 500μm (A, C, G), and 100μm (E).

To assess whether removing a single allele of *Ptch1* restores midface width in other transition zone mutants, we generated *Cc2d2a*^-/-^ *Ptch1*^-/+^ and *Cc2d2a*^-/-^ *Ptch1*^+/+^ embryos. As with *Tctn2*, removing a single allele of *Ptch1* restored midface width in *Cc2d2a* mutants (**Figure 5C,D**). Thus, reducing *Ptch1* gene dosage rescues midface expansion in both models of ciliopathy-associated hypoplasia.

As we had hypothesized that increased apoptosis underlay the midface hypoplasia of *Tctn2*^-/-^ *Ptch1*^+/+^ embryos, we assessed apoptosis in *Tctn2*^-/-^ *Ptch1*^+/-^ embryos via TUNEL staining. *Tctn2^-/-^ Ptch1^+/-^* embryos exhibited less apoptosis than *Tctn2*^-/-^ *Ptch1*^+/+^ embryos, with a restriction of apoptosis in the ventral telencephalon (**Figure 5E-F**). These results bolster the hypothesis that increased midline apoptosis accounts for the midline hypoplasia of transition zone mutants.

As genes encoding transition zone MKS components are epistatic to *Ptch1* (Reiter, 2006), we pondered how reducing *Ptch1* gene dosage restored facial midline development to transition zone MKS component mutants. The best studied role for PTCH1 is in repression of the HH signal transduction pathway. Therefore, we examined HH signal transduction pathway activity in *Tctn2^-/-^ Ptch1^+/+^* and *Tctn2^-/-^ Ptch1^+/-^* embryos. WM-ISH revealed that expression of neither *Shh* nor *Gli1* was increased in the ventral telencephalons of *Tctn2^-/-^ Ptch1^+/-^* embryos in comparison to *Tctn2^-/-^ Ptch1^+/+^* embryos (**Figure 5G**). Thus, the restoration of midface growth by reduction of *Ptch1* gene dosage is not due to a restoration of HH signal transduction.

In addition to its role in regulating HH signal transduction, PTCH1 exhibits pro-apoptotic activity in vitro (Thibert et al., 2003). As reducing *Ptch1* gene dosage reduces apoptosis without increasing HH signal transduction in *Tctn2* mutants, we conclude that it is the PTCH1-mediated death of the midline neurectorderm and facial ectoderm, and not alterations in HH signal transduction within those cells, that is the etiology of midface defects in transition zone mutants. Taken together, these results suggest a working model for how ciliary HH signaling regulates midface development.

In wild-type embryos, HH signaling within the prechordal plate is critical for *Gsc* expression and the induction of *Shh* in the adjacent neurectoderm and inhibition of apoptosis (**Figure 6A**). In transition zone mutants, defects in prechordal plate signaling cause reduced SHH in the neurectoderm, resulting in increased PTCH1-mediated cell death and midline collapse (**Figure 6B**). In transition zone mutants lacking a single allele of *Ptch1*, reduced SHH in the neurectoderm persists, but the attenuated PTCH1 is no longer sufficient to induce extensive cell death, allowing for normal midline facial development (**Figure 6C**).

**Figure 6.**
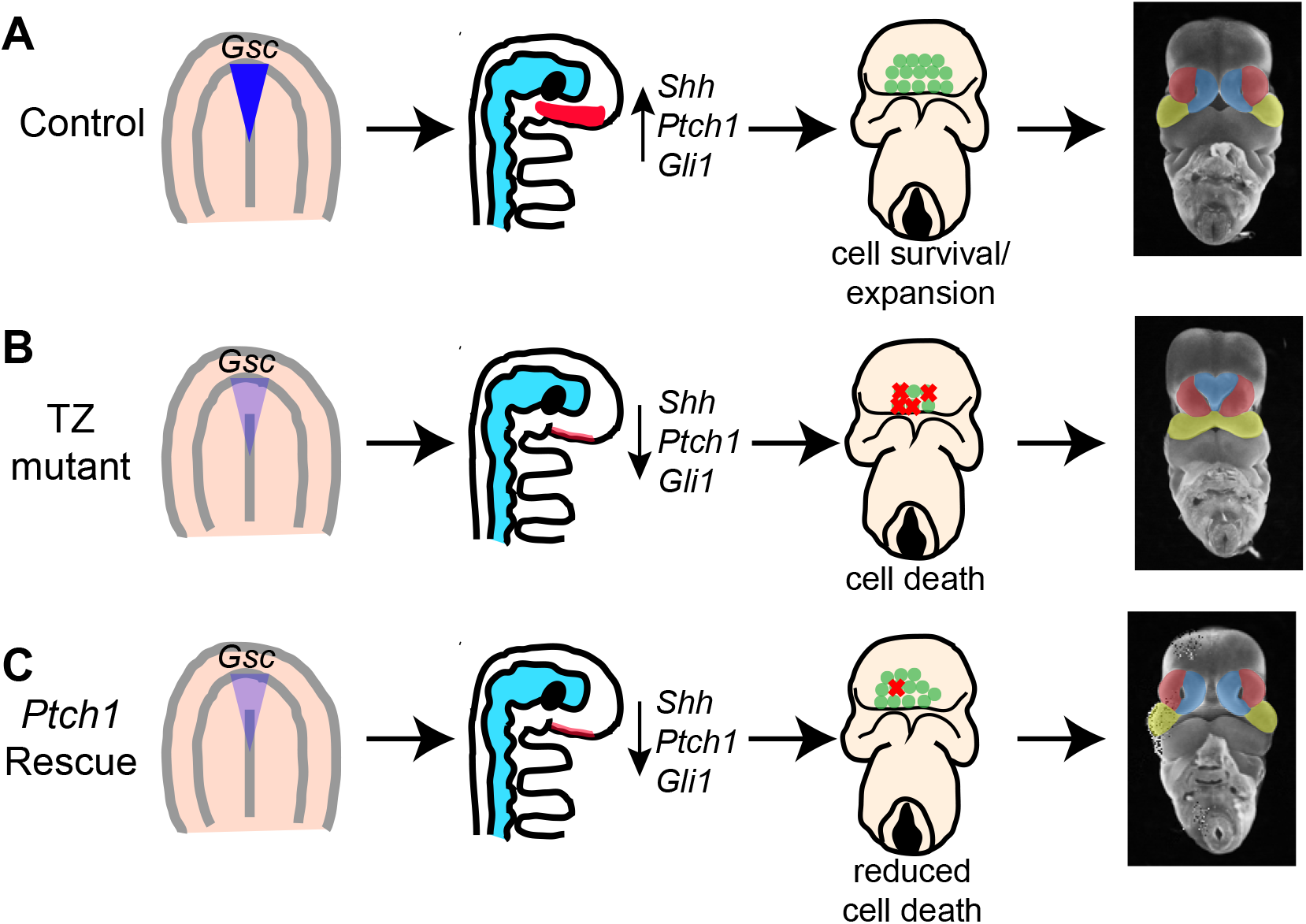
Model for transition zone coordination of facial midline development. (A) In wild-type embryos, the transition zone complex mediates signaling in the prechordal plate and HH pathway activation in the adjacent neurectoderm to allow for cell survival and normal midline development. (B) In transition zone mutants, disrupted signaling in the prechordal plate results in reduced *Shh* and HH pathway activation in the neurectoderm, resulting in increased cell death and corresponding collapse of the facial midline. (C) In transition zone mutants with *Ptch1* haploinsufficiency (*Ptch1* rescue), reduced cell death allows for normal midface development despite persistant reduction of *Shh* and HH pathway activation in the neurectoderm.

## Discussion

Using a combination of genetic, developmental and biochemical techniques, we have identified a common etiology by which disruption of MKS transition zone proteins (TCTN2, CC2D2A, and TMEM231) result in midline hypoplasia and hypotelorism. We traced the origin of the molecular defect contributing to the midline phenotype to the prechordal plate, defects in which resulted in reduced HH pathway activation and cell survival in the adjacent neurectoderm and facial midline collapse. We uncovered *Ptch1* gene dosage as a key mediator of cell survival in the facial midline of transition zone mutants, as loss of a single allele of *Ptch1* rescued cell survival and midline development in *Tctn2* mutants. Together, these results reveal a new paradigm whereby primary cilia mediate signal crosstalk from the prechordal plate to the adjacent neurectoderm to promote cell survival, without which the facial midline collapses and hypotelorism results.

Different ciliopathies are associated with either narrowing or expansion of the facial midline (hypotelorism or hypertelorism) (Schock et al., 2015; Zaghloul & Brugmann, 2011). Severe ciliopathies associated with perinatal lethality, such as Meckel syndrome, can present with hypotelorism or hypertelorism while other ciliopathies such as Joubert syndrome typically present with hypertelorism (Brugmann, Cordero, & Helms, 2010b; MacRae, Howard, Albert, & Hsia, 1972; Schock & Brugmann, 2017). How disruption in primary cilia can give rise to these opposing phenotypes has been an active area of interest. Hypertelorism is attributable to roles for cilia in promoting Gli3 repressor formation in neural crest cells (Brugmann, Allen, James, Mekonnen, et al., 2010a; C. F. Chang et al., 2014; C.-F. Chang, Chang, Millington, & Brugmann, 2016; Liu, Chen, Johnson, & Helms, 2014). Our work implicates a distinct etiology of hypoterlorism: rather than involving neural crest, midline hypoplasia can be caused by defects in the ciliary transition zone in the prechordal plate.

The earliest alteration we detected in *Tctn2*^-/-^ signaling centers that regulate craniofacial development was in the prechordal plate at the end of gastrulation. TCTN2 was dispensable for the expression of *Shh* and *T* in the prechordal plate, indicating that induction and specification of the prechordal plate were unaffected. In contrast, TCTN2 was essential for prechordal plate expression of *Gli1* and *Gsc*. In tissues such as the limb bud, TCTN2 is dispensable for ciliogenesis but critical for ciliary HH signaling and the induction of HH target genes such as *Gli1* (Ben Chih et al., 2011; Dowdle et al., 2011; Garcia-Gonzalo et al., 2011). We found that, similarly in the prechordal plate, TCTN2 is dispensable for ciliogenesis but critical for induction of *Gli1*.

In many developmental events, such as limb patterning, SHH signals to neighboring cells to induce a pattern (Panman & Zeller, 2003; Zhulyn et al., 2014). In other developmental events, such as notochord to neural tube signaling, SHH signals produced by the notochord induce the expression of *Shh* in the overlaying neural tube (Fuccillo et al., 2006). SHH produced by the prechordal plate may fall into the latter category, as the absence of *Shh* expression in *Tctn2*^-/-^ mutants presages reduced *Shh* expression and reduced expression of HH pathway transcriptional targets *Gli1* and *Ptch1* in the region of the basal forebrain sometimes referred to as the rostral diencephalon ventral midline. Thus, in the posterior midline, the notochord induces *Shh* expression in overlying neuroectoderm, and in the anterior midline, the prechordal plate induces *Shh* expression in the overlying neuroectoderm. In the failure of the prechordal plate to induce Shh expression in the overlying neurectoderm, *Tctn2* mutants recapitulate previous observations of *Lrp2* mutants which display attenuated responses to SHH (Christ et al., 2012). One possible mechanism by which SHH may activate expression of *Shh* in the basal forebrain is via the induction of the transcription factor SIX3. SIX3 is regulated by HH signaling and required for the induction of *Shh* in the developing forebrain (Geng et al., 2008; Jeong et al., 2008). However, our observation that *Six3* expression is unaltered in in the forebrains of *Tctn2* mutants diminishes support for this hypothesis.

In caudal neural tube and limb patterning, HH signals induce patterning. In hair follicles and cerebellar granule cells, HH signaling promote proliferation. In addition to roles in patterning and proliferation, HH signals can bind to PTCH1 to inhibit apoptosis (Aoto & Trainor, 2015; Borycki et al., 1999; Thibert et al., 2003). Our data are consistent with a role of PTCH1 in promoting apoptosis in the neuroectoderm and facial ectoderm which is inhibited by prechordal plate-produced SHH. In the absence of TCTN2, the prechordal plate does not produce SHH, releasing PTCH1 to promote apoptosis in the midline and resulting in midface hypoplasia. This model is consistent with previous data demonstrating that surgical ablation of the prechordal plate reduces the forebrain (Aoto et al., 2009). Increased cleaved Caspase-3 and Caspase-9 staining in the basal forebrain and facial ectoderm of E9.5 *Tctn2* mutants provides further support for apoptosis contributing to the midline defect.

We sought to identify where TZ function is critical to coordinate normal midline facial development by taking a tissue-specific approach to delete *Tctn2* in the various tissues that comprise the craniofacial complex. Deletion of *Tctn2* in the prechordal plate (via *Isl1-cre*) or the neurectoderm (via *Sox1-cre*) did not recapitulate the midline hypoplasia seen in *Tctn2* global null mutants (Supplemental figure 3). Similarly, deletion of *Tctn2* in the facial ectoderm (via *Crect-cre*) or in the forebrain and facial ectoderm (via *Foxg1-cre*) also failed to result in midline collapse (Supplemental figure 4). This result could be due to residual TCTN2 function at these early timepoints as indicated by persistent ARL13B localization at the cilium, low turnover of TZ components, and an inability to delete *Tctn2* at an early developmental timepoint where the TZ coordinates midline facial development. New genetic tools that will allow an earlier deletion of *Tctn2* in craniofacial complex tissues will be critical to uncover the tissue-specific requirements of *Tctn2* in midline development.

Surprisingly, removing one allele of *Ptch1* fully rescues the midface defect in both *Tctn2* and *Cc2d2a* transition zone cilia mutants. Even more surprisingly, this phenotypic rescue is not associated with restoration of either *Shh* expression or HH pathway activation in the basal forebrain. We propose that reducing *Ptch1* levels attenuates the PTCH1-mediated pro-apoptotic program normally attenuated by SHH in the basal forebrain.

In summary, we have identified the primary cilia transition zone as a critical regulator of facial midline development. The transition zone component TCTN2 was critical for SHH signaling in the prechordal plate and uncovered a signaling paradigm whereby the transition zone promotes cell survival by mediating crosstalk between the prechordal plate and neurectoderm to promote HH pathway activation. These results provide new insights into how primary cilia mediate cell survival to promote facial development.

## Materials and Methods

### Mouse Strains

All mouse protocols were approved by the Institutional Animal Care and Use Committee (IACUC) at the University of California, San Francisco. *Tctn2^+/-^* (*Tctn2^tm1.1Reit^), Cc2d2a^+/-^ (Cc2d2a^Gt(AA0274)Wtsi^), Tmem231^+/-^ (Tmem231^Gt(OST335874)Lex^), and Tmem67^+/-^ (Tmem67^tm1Dgen^)* mouse alleles have been previously described. *Wnt1-cre* (*Tg(Wnt1-cre)11Rth)* and *Islet1-cre (Isl1^tm1(cre)Sev^)* mice were obtained from Brian Black, *Sox1-cre* (*Sox1^tm1(cre)Take^)* mice were obtained from Jeff Bush, *Foxg1-cre* (*Foxg1^tm1(cre)Skm^)* mice were obtained from Stavros Lomvardas, and *Crect-cre* (*Tg(Tcfap2a-cre)1Will)* mice were obtained from Trevor Williams. The *Ptch1^tm1Mps^* allele was used in this study as a null allele. All mice were maintained on a C57BL/6J background. For timed matings, noon on the day a copulation plug was detected was considered to be 0.5 days postcoitus.

### Immunofluorescence

The antibodies used in this study were rabbit α-Arl13b (1:1000, Proteintech 17711-1-AP), rabbit α-cleaved-caspase 3 (Asp175) (1:400, Cell Signaling #9664), rabbit α-cleaved-caspase 9 (Asp353) (1:100, Cell Signaling #9509), rabbit α-Phospho-Histone H3 (Ser28) (1:400, Cell Signaling #9713), chicken anti-GFP (1:1000, Aves labs GFP-1020), goat gamma-tubulin (1:200, Santa Cruz sc7396), rat E-Cadherin (1:1000, Invitrogen 13-1900), and rabbit FoxA2 (1:400 abcam ab108422). The In Situ Cell Death Detection Kit, Fluorescein (Roche) was used for TUNEL cell death assay. For immunofluorescence antibody staining of frozen tissue sections, embryos were fixed overnight in 4% PFA/PBS, washed in PBS and cryopreserved via overnight incubation in 30% sucrose/PBS. Embryos were embedded in OCT and frozen at -80C. Frozen OCT blocks were cut into 10μM sections. For immunostaining, frozen sections were washed 3×5’ in PBST (0.1%Tween-20/PBS) followed by blocking for 2 hours in blocking solution (5% donkey serum in PBS+ 0.3% Triton X-100 + 0.2% Na-azide). Slides were incubated overnight in primary antibody diluted in blocking solution at 4 degrees. The following day, slides were washed 3×10’ in PBST, stained with appropriate AlexaFluor 488, 568, or 647 conjugated secondary antibodies (Life Technologies) at 1:1000 and Hoecsht or Dapi nuclear stain in blocking buffer for 1 hour, rinsed 3×10’ with PBST and mounted using Fluoromount-G (Southern Biotech). All steps performed at room temperature unless otherwise noted. *Note: For gamma-tubulin antibody staining, antigen retrieval by incubating with 1%SDS/PBST for 5 mins prior to blocking and primary antibody incubation in required for good staining. Stained samples were imaged on a Leica SP-5 confocal microscope. Images were processed using FIJI (ImageJ).

### In Situ Hybridization

Whole mount in situ hybridization was performed as previously described (Harrelson, Kaestner, & Evans, 2012). DIG-labeled riboprobes were made using plasmids from the following sources: *Shh* (Echelard et al., 1993), *Gsc* (Blum et al., 1992), *Ptch1* (Goodrich, 1999), *Foxa2* (Brennan et al., 2001), *Gli1* (EST W65013).

### RT-qPCR

For gene expression studies, RNA was extracted from E8.5/E9.5 embryo heads using an RNAeasy Micro Kit (QIAGEN), and cDNA synthesis was performed using the iScript cDNA synthesis kit (BioRad). RT-qPCR was performed using EXPRESS Sybr GreenER 2X master mix with ROX (Invitrogen) and primers homologous to *Shh, Ptch1, Gli1,* or *Six3* on an ABI 7900HT real-time PCR machine. Expression levels were normalized to the geometric mean of three control genes (*Actb, Hprt,* and *Ubc)*, average normalized Ct values for control and experimental groups determined and relative expression levels determined by Δ ΔCt. The RT-qPCR of each RNA sample was performed in quadruplicate and each experiment was replicated.

### Embryo processing for midface imaging

Embryos were harvested in ice-cold PBS, staged by counting somite number, and fixed o/n at 4 degrees in 4%PFA/PBS. Embryo heads were removed and stained in 0.01% ethidium bromide in PBS at room temperature for 15 minutes. Embryos were positioned using glass beads in PBS and imaged on a Leica MZ16 F fluorescence stereomicroscope.

### Image Quantification

For 2D midface width quantification the infranasal distance was measure using FIJI software by drawing a line between the center of each nasal pit. For quantification of cell death and proliferation, 6 sections per embryo were quantified. Staining with epithelial marker E-cadherin was used for quantification of facial ectoderm while nuclear morphology was used to separate mesenchyme, hindbrain and forebrain tissue compartments. For quantification, threshold was first set for each image followed by binary watershed separation to obtain accurate nuclei counts. The percentage of TUNEL+ or pHH3+ nuclei were compared between *Tctn2* mutant and control samples.

---

**Table 1.**
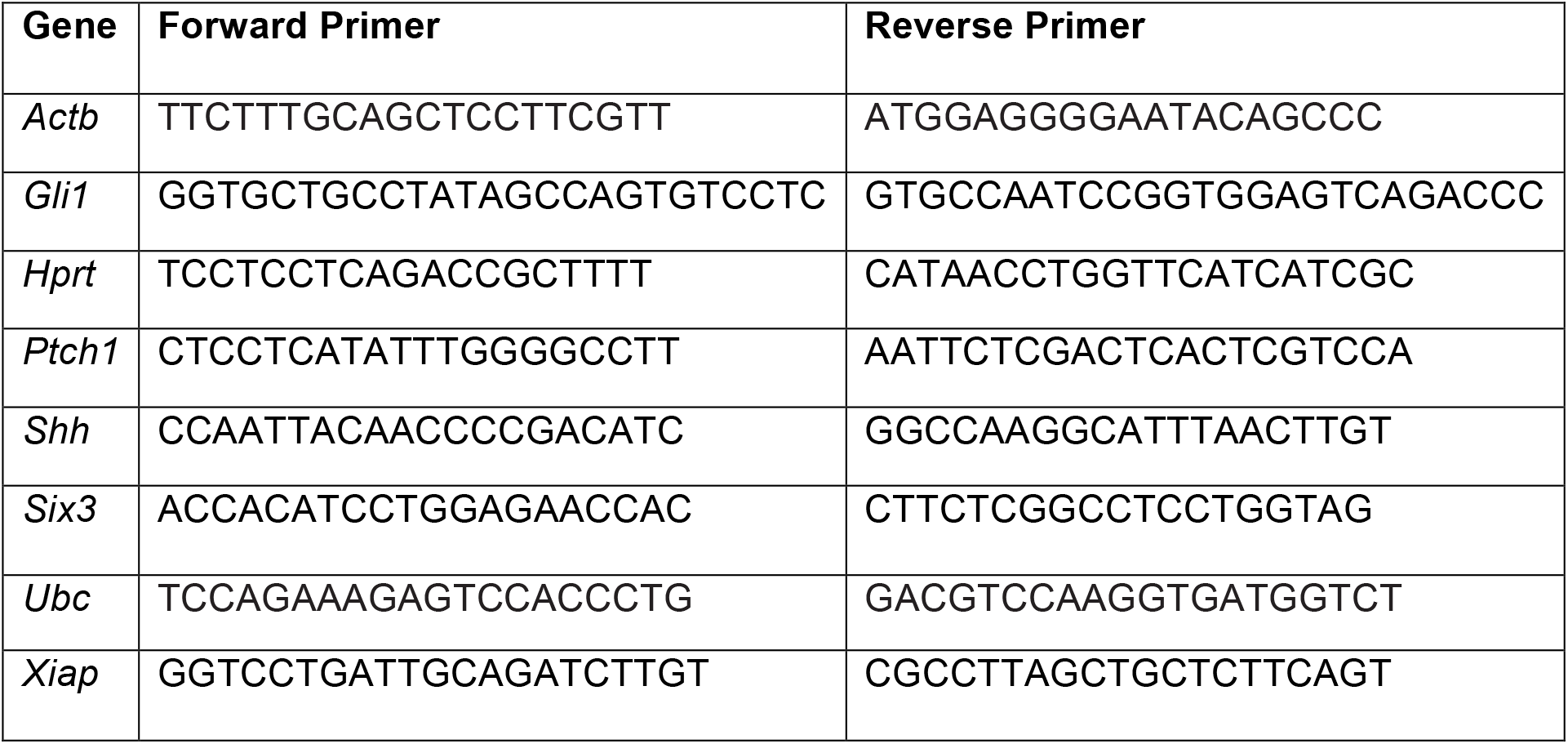
Primer sets used for RT-qPCR.

**Table 2.**
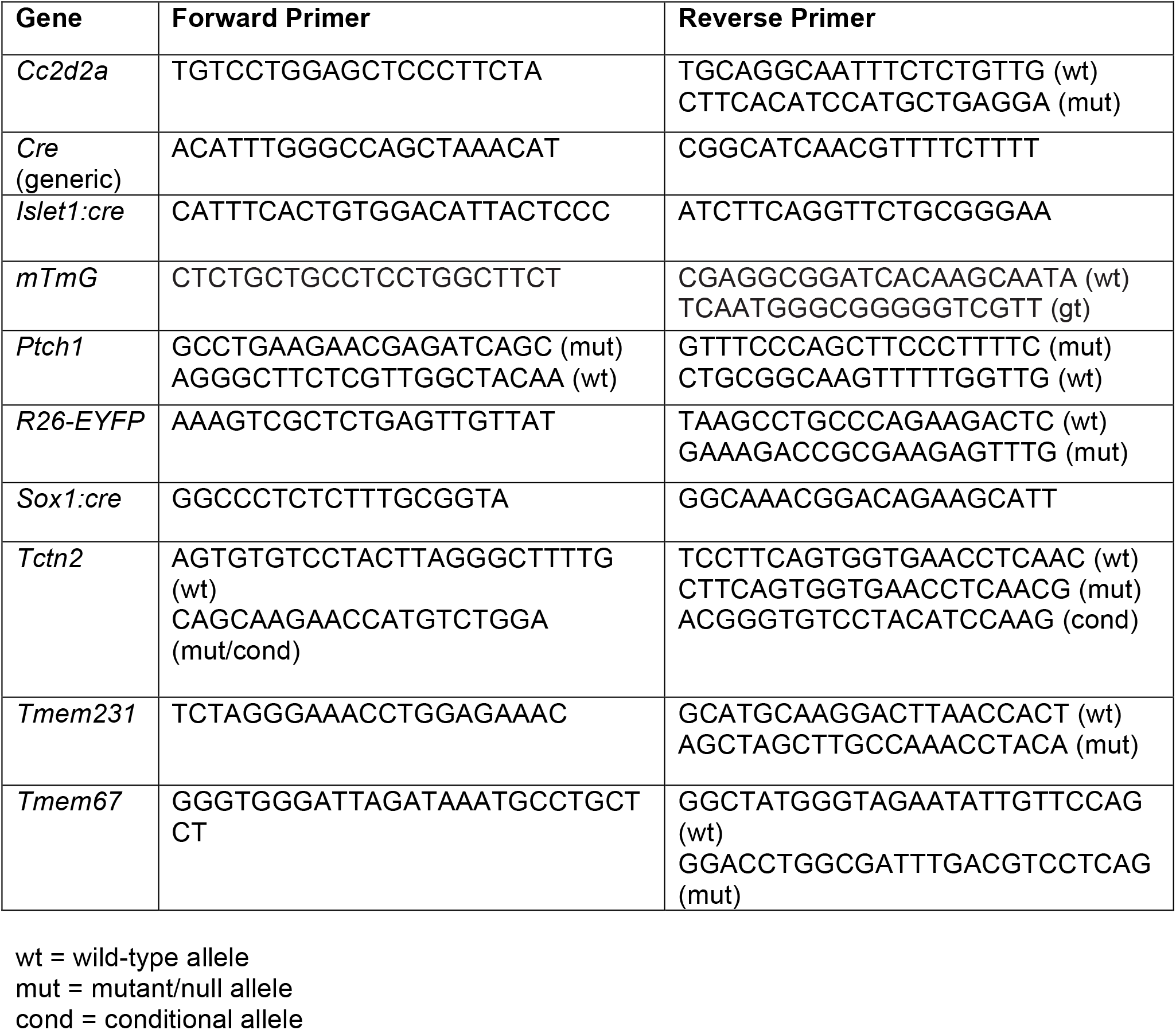
Primer sets used for Genotyping Assays.

## Supplemental Materials

**Supplemental Figure 1.**
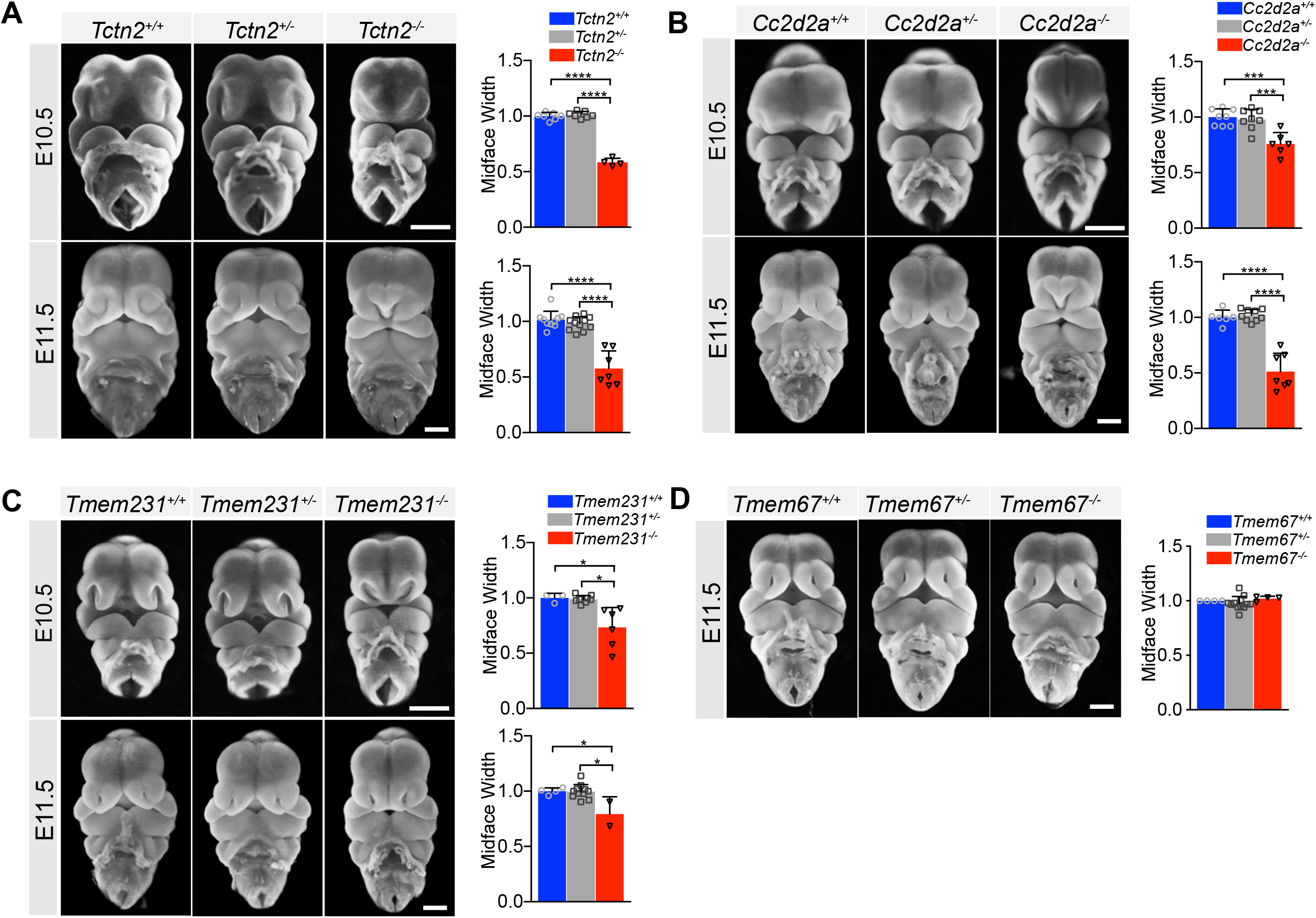
The ciliary MKS transition zone complex is essential for midline facial development. Reproduced frontal view images from Figure 1 of *Tctn2* (A), *Cc2d2a* (B), *Tmem231* (C), wildtype and null embryos at E10.5 and E11.5 with the addition of heterozygous embryos. (D) *Tmem67* null and wildtype E11.5 embryos reproduced from Figure 1 with addition of heterozygous embryos. Quantification of midface width at respective timepoints measured via one-way Anova followed by Tukey’s multiple comparisons test. Error bars represent the standard deviation (SD). Scale bar indicates 500μm. p<0.05 (*), p<0.0002 (***), p<0.0001(****).

**Supplemental Figure 2.**
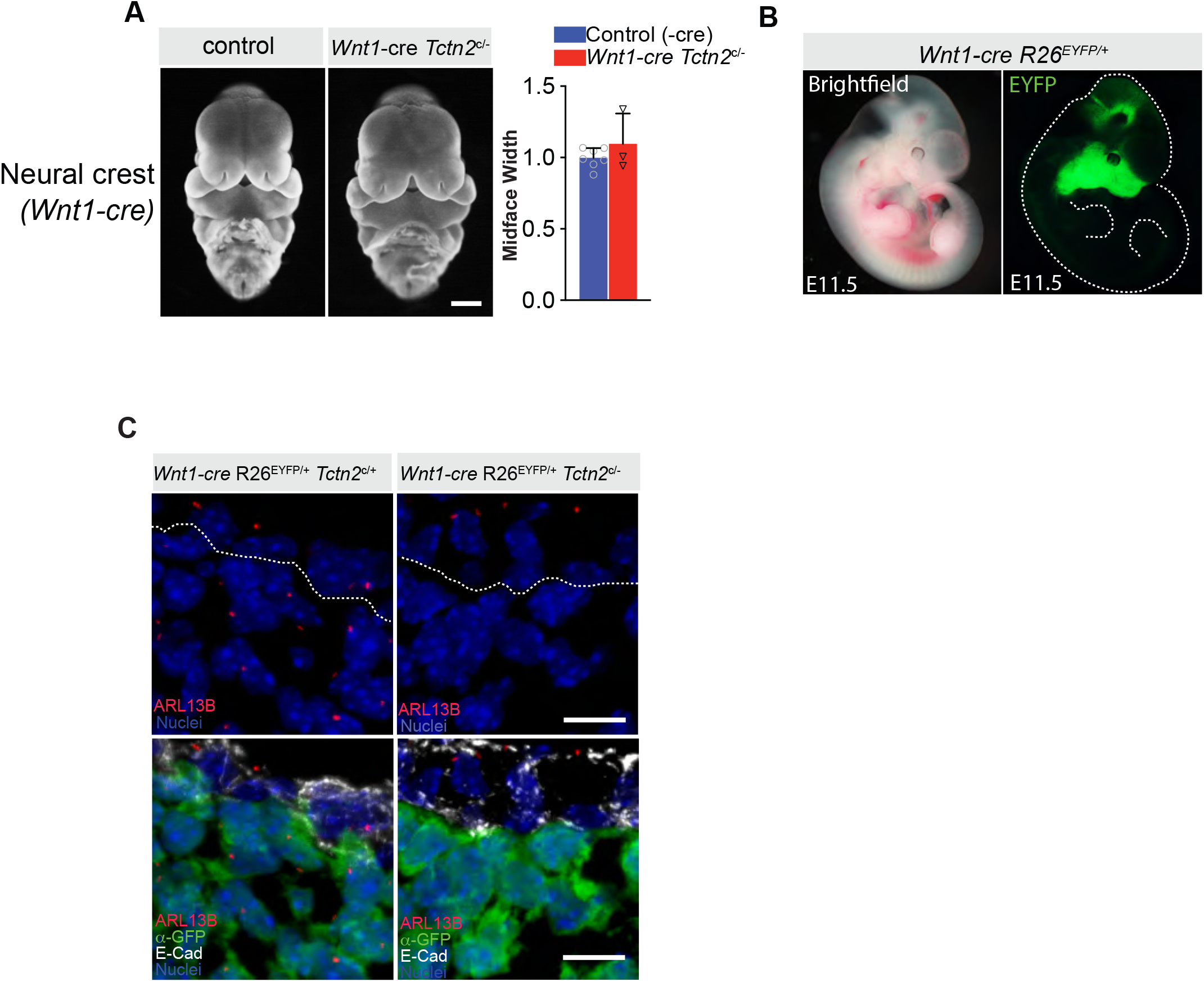
Deletion of *Tctn2* in the neural crest using the *Wnt1-cre* driver does not result in hypotelorism. (A) Frontal view images of E11.5 *Tctn2* control (*Tctn2*^c/-^) and neural crest deletion (*Wnt1-cre Tctn2*^c/-^) embryos with corresponding midface width quantification. (B) Brightfield and fluorescent images of E11.5 *Wnt1-cre R26*^EYFP/+^ embryos show expected pattern of cre recombination (embryo outlined with white dotted line in B). (C) Immunofluorescence staining of E10.5 medial nasal prominence sections for ciliary membrane protein ARL13B, GFP, and epithelial marker E-Cadherin (E-Cad). Dotted white line in (C) represents boundary separating epithelia and neural crest mesenchyme. Scale bars indicate 500μm (A), and 10μm (C).

**Supplemental figure 3.**
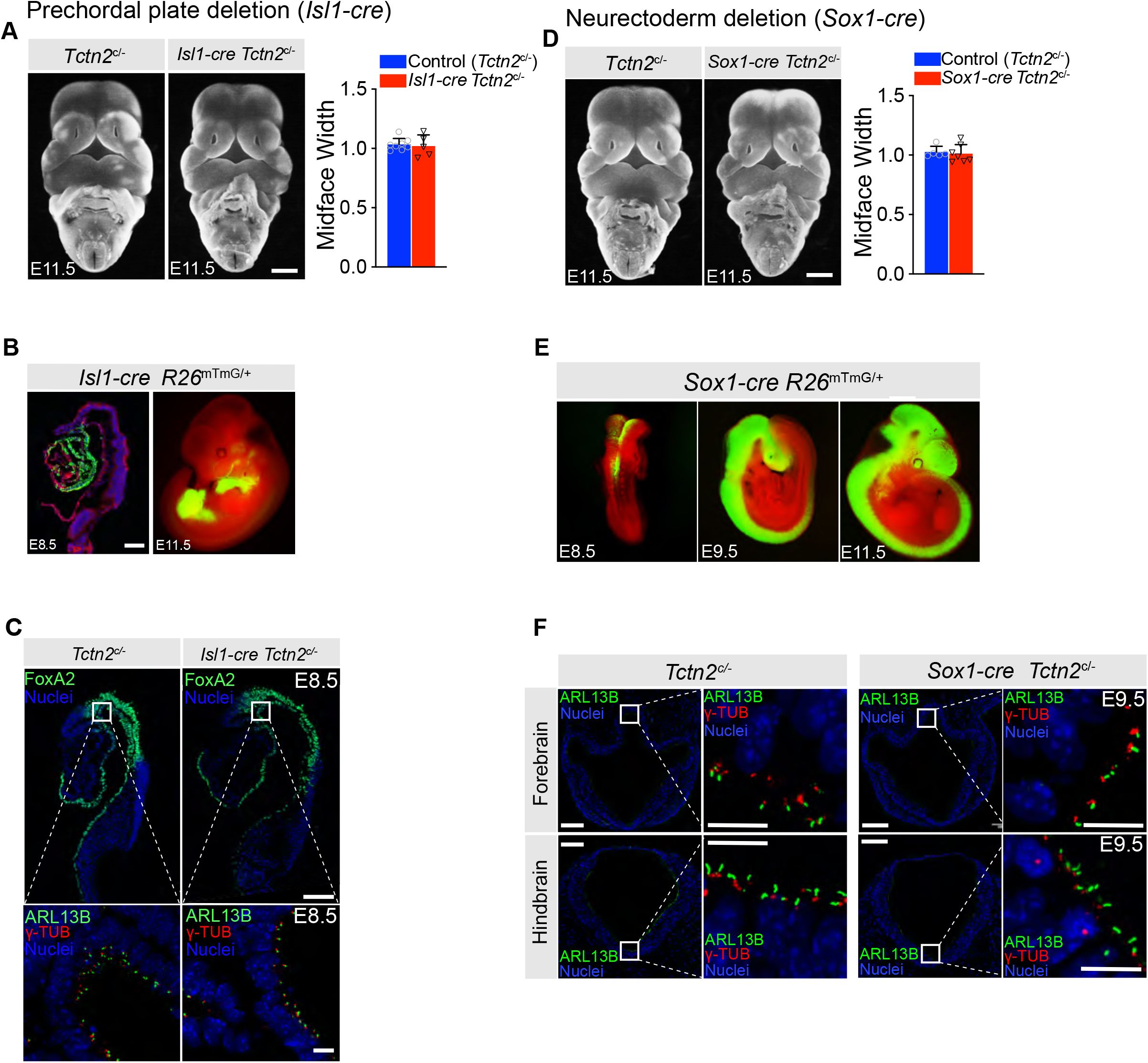
Deletion of *Tctn2* in the prechordal plate via the *Isl-1cre* driver or neurectoderm via the *Sox1-cre* driver does not result in hypotelorism. (A) Frontal view images of E11.5 control (*Tctn2*^c/-^) and mutant (*Isl1-cre Tctn2*^c/-^) embryos with corresponding midface width quantification. (B) Analysis of *Isl1-cre* recombination pattern at E8.5 and E11.5 using the *R26*^mTmG^ reporter shows robust recombination in the prechordal plate and endodermal derivatives at respective timepoints. (C) Analysis of *Tctn2* deletion in the prechordal plate of E8.5 embryos through ciliary localization of ARL13B indicates residual TCTN2 function remains at this early time point despite reporter recombination. (D) Frontal view images of E11.5 control (*Tctn2*^c/-^) and neurectoderm-specific *Tctn2* mutant (*Sox1-cre Tctn2*^c/-^) embryos with corresponding midface width quantification. (E) Analysis of *Sox1-cre* recombination pattern at E8.5, E9.5 and E11.5 using the *R26*^mTmG^ reporter shows early recombination in the neural folds at E8.5 followed by robust recombination throughout the neurectoderm at E9.5 and E11.5. (F) Analysis of *Tctn2* deletion in the neurectoderm of E9.5 embryos through ciliary localization of ARL13B in the forebrain (top panels) and hindbrain (bottom panels) indicates residual TCTN2 function remains at this early time point despite robust reporter recombination. Scale bars in A and D indicates 500μm, in B and low-magnification images in C and F indicates 100μm, and in high-magnification images in C (bottom panels) and F (right panels) indicates 10μm.

**Supplemental figure 4.**
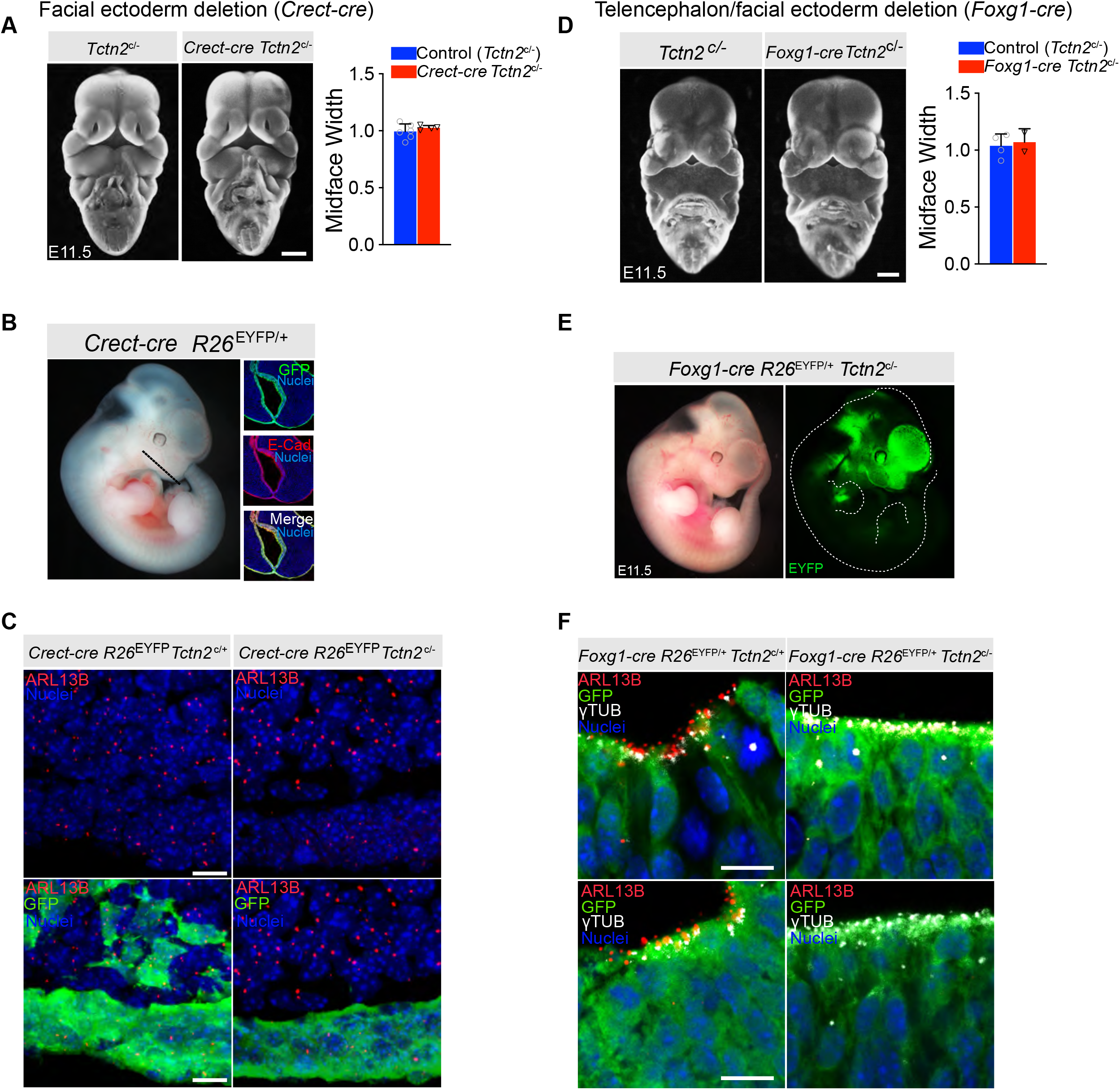
Deletion of *Tctn2* in the facial ectoderm via the *Crect-cre* driver or forebrain and facial ectoderm via the *Foxg1-cre* driver does not result in hypotelorism. (A) Frontal view images of E11.5 control (*Tctn2*^c/-^) and facial ectoderm deletion (*Crect-cre Tctn2*^c/-^) embryos with corresponding midface width quantification. (B) Analysis of *Crect-cre* recombination pattern at E11.5 using the *R26*^EYFP^ reporter shows robust recombination in the facial ectoderm as evidenced by co-localization with epithelial marker E-Cadherin (E-Cad). (C) Analysis of *Tctn2* deletion in the facial ectoderm of E11.5 embryos through ciliary localization of ARL13B indicates residual TCTN2 function remains at this time point as ARL13B accumulation in cilia persists in the facial ectoderm of *Crect-cre Tctn2*^c/-^ mutants. (D) Frontal view images of E11.5 control (*Tctn2*^c/-^) and forebrain/facial ectoderm-specific *Tctn2* deletion (*Foxg1-cre Tctn2*^c/-^) embryos with corresponding midface width quantification. (E) Analysis of *Foxg1-cre* recombination pattern at E11.5 using the *R26*^EYFP^ reporter shows expected robust recombination in the forebrain and facial ectoderm at E11.5. (F) Immunofluorescence staining of E11.5 neural tube sections for ciliary membrane protein ARL13B, basal body marker gamma tubulin (γTUB), and α-GFP for readout of reporter recombination shows loss of TCTN2 function in this tissue through loss of ARL13B ciliary accumulation in mutant embryos. Dotted line in B indicates approximate section through midface for immunofluorescence images in B. Dotted white line in E indicates embryo outline.

## Notes

### Competing Interest Statement

The authors have declared no competing interest.

